# Composite mutations give an extra insight into epistasis

**DOI:** 10.1101/2022.06.16.496391

**Authors:** Evgenii M. Zorin, Carolina M. Erazo, Dmitry N. Ivankov

## Abstract

The intricate genotype-phenotype relationship has been a long-standing issue in biology, important both from the fundamental and applied points of view. One of the major irregularities hindering progress in establishing these links is epistasis – the complex and elusive interaction between mutations. Despite the vast accumulated genetic data and progress in this area, epistasis is still far from being completely understood. Epistasis can be studied quantitatively in combinatorially complete datasets, which form hypercubes in protein sequence space, where connected sequences are one mutation away from each other. However, this might be insufficient to portray the full picture of epistatic interactions. To extend the repertoire of the methods for exploring epistasis, we propose here to consider hyperrectangles, where some edges connect sequences being two or more mutations away from each other. The present work formalizes the theoretical knowledge about these novel structures and compares the amount of epistasis identified in hypercubes and hyperrectangles constructed from experimental datasets. A new algorithm, CuboidME, was developed for calculating hyperrectangles, which were then compared to hypercubes. In the experimental datasets, there were four orders of magnitude more hyperrectangles than hypercubes for the same sample size. Subsequently, we showed that for the studied datasets there is an increase in epistasis measured by epistatic coefficients in hyperrectangles compared to hypercubes. For the same datasets, hyperrectangles could find more sign epistasis than using hypercubes alone. We also show that there is a trend for increase in epistasis with increasing number of mutations being considered in a hyperrectangle. The results indicate that hyperrectangles can be used to reveal more information on epistasis in a fitness landscape, especially if it is combinatorially incomplete.

## 1. Introduction

Genotype-phenotype relationship is one of the central issues in evolutionary, developmental, and systems biology (de Visser et al., 2011). Understanding this relationship is critical in predicting disease from genotype and development of better vaccines and antibiotics through our enhanced understanding of microorganismal evolution. Phenotype, or observable traits of an organism, determines both the susceptibility to certain genetic diseases and adaptability to survive in the environment in question. Among the factors that largely determine phenotype, genotype is the principal heritable factor; therefore, understanding how genotype affects phenotype will allow us to predict evolutionary processes and determine which phenotypic characteristic an organism would have. Namely, an enhanced understanding of genotype’s effect on phenotype leads to numerous applications in the fundamental research and practical fields, such as predicting risk of disease from genotype, personalized medicine, developing vaccines and antibiotics through our enhanced understanding of microorganismal evolution, and control of crop pests (Kemble et al., 2019; Nosil et al., 2020). Yet, this relationship is far from being linear due to epistasis – the deviation in fitness from the additive effects of individual mutations (Breen *et al*., 2012; Sarkisyan *et al*., 2016; Weinreich *et al*., 2013; Phillips, 2008) – making the investigation of epistasis an important task. The linkage of fitness to genotype space can be viewed as an abstraction in the fitness landscape (Wright, 1932). The most straightforward way to study epistasis are combinatorially complete datasets, forming hypercubes in genotype space (Weinreich *et al*., 2013; Poelwijk et *al*., 2016; Esteban et *al*., 2019). The value of combinatorially complete datasets for the study of evolution and epistasis was shown convincingly for a number of evolutionary landscapes (Weinreich *et al*., 2013).

Previously, we published a recursive algorithm for finding all hypercubes in fitness landscape experiments that use quasi-random mutagenesis, which would be infinitely slow for a brute-force approach (Esteban et al., 2019). Using the algorithm should be useful for investigating high-order epistasis in the high-throughput experimental data (Esteban et al., 2019; Erazo et al., manuscript in preparation). From an evolutionary point of view, we are reduced to studying single evolutionary effects, most of which being single nucleotide polymorphisms (SNPs) comprising hypercubes. However, using hypercubes gives only information about local neighborhood in the fitness landscapes. Moreover, for sparse landscapes the hypercubes may not be identified at all.

Here, we propose consideration of composite mutations consisting of multiple single mutations. We generalize analysis of the combinatorially complete dataset using composite mutations. We argue that composite mutations allow finding many more epistatic characterizations of fitness landscapes than by using hypercubes. We show that, with the same amount of information obtained for hypercubes, the data obtained from the viewpoint of composite mutations contain much more information about epistasis and allow a more insightful glance into the nature of epistatic interactions.

## 2. Methods

### 2.1. Fitness landscape datasets

We took for our analysis local fitness landscapes of green fluorescent protein (GFP) (Sarkisyan et al., 2016), segments 1-12 (S1-S12) of IGPD (imidazoleglycerol-phosphate dehydratase) (Pokusaeva et al., 2019), RNA recognition motif (RRM2) of *Saccharomyces cerevisiae* poly(A)-binding protein (Melamed et al., 2013), and WW domain of hYAP65 protein (Araya et al., 2012).

All fitness landscapes were converted into the format used in (Esteban et al., 2019): 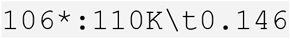, where position is followed by the amino acid residue substituting the residue in the reference sequence (asterisk denoting stop codon).

### 2.2. Algorithm

The algorithm was written using Python 3.9.6 in Visual Studio Code IDE. The detailed description of the principle of work and notation of hyperrectangles is described in the subsection 3.1 of the Results section.

### 2.3. Calculating epistatic coefficients

In the present research work, we analyze pairwise epistasis expressed in coefficients *a*_12_ from a thermodynamic matrix for 2D hypercubes; therefore, epistatic coefficients were found using triangular transform matrix: (Poelwijk *et al*., 2016):

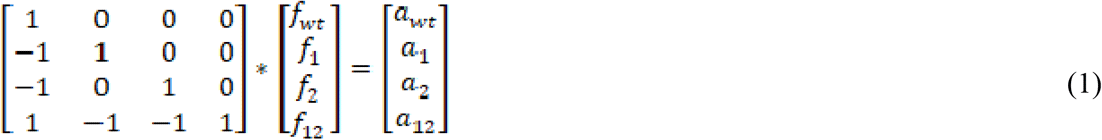

We chose the thermodynamic matrix described by (Poelwijk et al., 2016), as it was the most straightforward one and easier to understand. For the purposes of the present work, the choice of a matrix for calculating epistatic coefficients is not imperative, as hypercubes and hyperrectangles were compared based on the same method of calculating epistasis. The performance of different matrices for obtaining epistatic coefficients is a task for a separate research project and was not considered in this article.

### 2.4. Statistical analysis

Distributions of epistatic coefficients *a*_12_ were checked for normality by using R to construct Q-Q plots and carry out the Anderson-Darling normality tests. Subsequently, the choice of statistical test was either an unpaired t-test (in case of normal distribution) or Mann-Whitney U test (in case of non-normal distribution). Mann-Whitney tested the following hypotheses:

- H_0_: there is no difference between the distributions of epistatic coefficients derived from hypercubes and hyperrectangles;
- H_A_: there is a difference between the two distributions (two-tailed test); mean of distribution of hyperrectangles is greater than that of hypercubes (one-tailed test).

The choice of one-tailed or two-tailed test is specified in each chapter applying the statistical test.

## 3. Results

### 3.1. Hyperrectangles as a novel structure in sequence space

For the purpose of the analysis, we introduce two new terms – hyperrectangles and hypercuboids. Hyperrectangles represent combinatorially complete datasets including necessary edges representing composite mutations (Figure 1). Hypercuboids comprise both hypercubes and hyperrectangles (Figure 1). In the present work, we study hypercuboids for amino acid sequences; however, the same analysis could be performed on nucleotide sequences as well. Figure 1 provides an example of 2D hypercube and hyperrectangle regardless of the other structures present in a combinatorially complete dataset; Figure 2 provides an example of 2D hypercube and hyperrectangle in the context of 3D hypercube.

**Figure 1.**
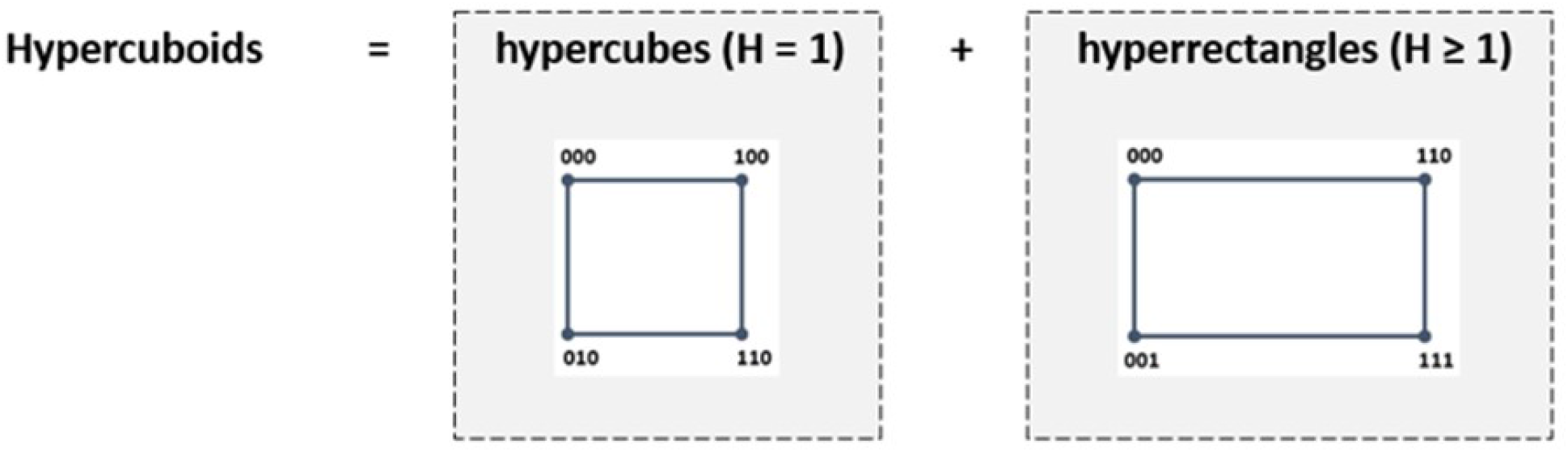
A visual explanation of the related terms hypercubes, hyperrectangles, and hypercuboids (H – hamming distance; amount of mutations separating two genotypes). Hypercuboids comprise hypercubes, in which each genotype connects to other genotypes if they are H = 1 mutation away, and hyperrectangles, in which some genotypes are connected if they are H = 2 and more mutations away from each other.

**Figure 2.**
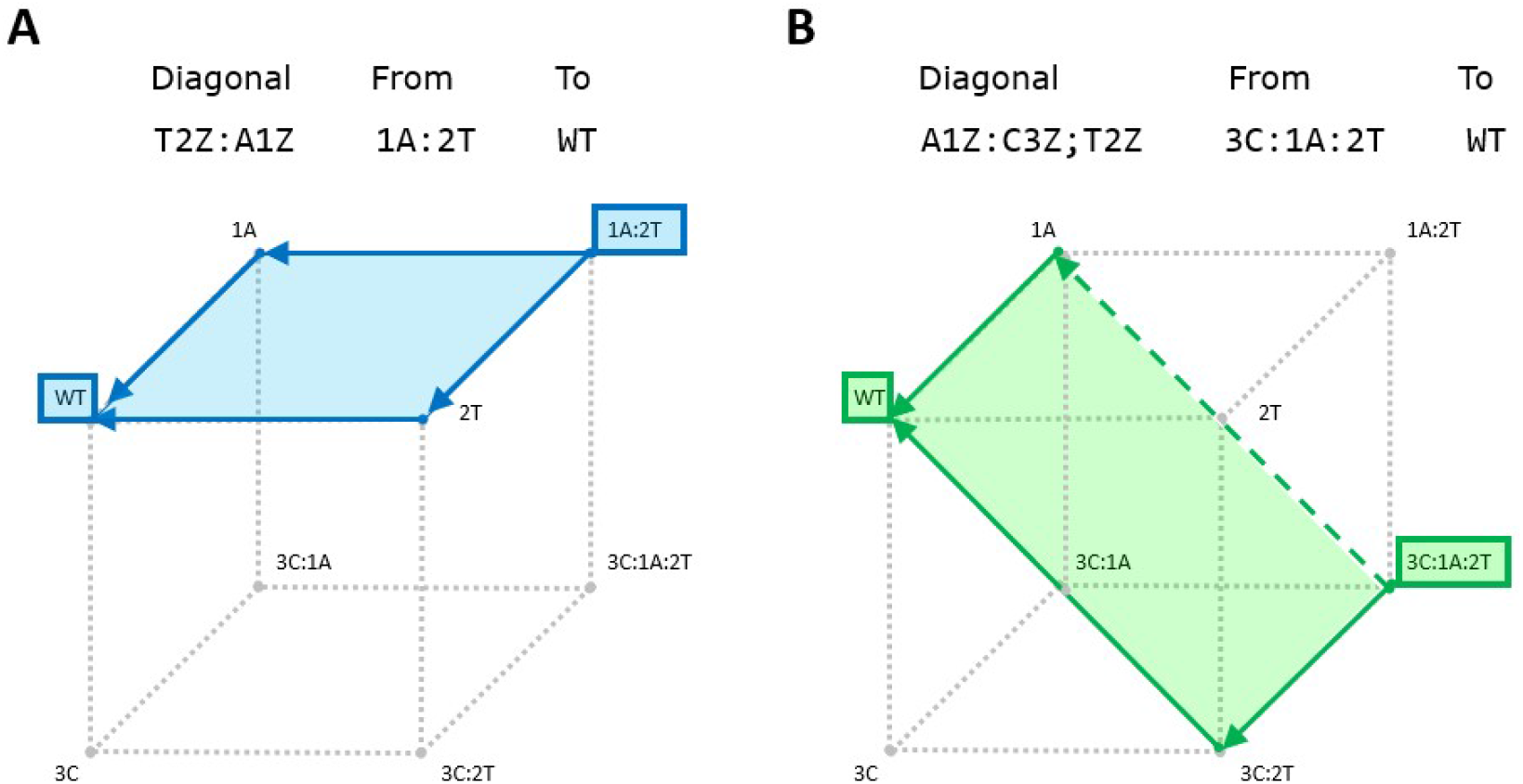
Examples of 2D hypercube and hyperrectangle in a combinatorially complete dataset of K = 3 mutational positions. Mutational pathways are given as leading from one sequence to another that is the closest to the WT variant. Sequences are represented as mutations added to the WT (e.g. 3C, mutation in position 3 to amino acid C). Letters A, T, C denote amino acids; letter Z – residue variant belonging to the WT sequence. **A**) A hypercube with WT, full mutant with two mutations 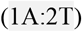 and a diagonal representing the mutational pathway from the full mutant to the WT variant; **B**) A hyperrectangle containing a diagonal with one point mutation 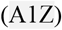 and a composite mutation 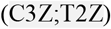.

The notation of hypercubes is similar to that given in (Esteban et al., 2019). In brief, for a hypercube of any dimension we are given genotypes – WT, full mutant, and all genotypes in between. The key concept when considering both hypercubes and hyperrectangle is a diagonal which connects most distant genotypes in a hyperstructure. To denote diagonal in written form for hypercubes, mutations connecting the two genotypes are separated by a colon (Esteban *et al*., 2019) (Figure 2A). The notation for hyperrectangles builds upon the previous one, with an addition of a semicolon, which separates individual mutations within a composite mutation (Figure 2B).

The first step is generating 1D hyperrectangles from the sequences given in Table 1, which is accomplished by performing a pairwise comparison of all sequences. For each sequence pair a diagonal is generated, which consists of mutations needed to get from one sequence to the other. By default, all mutations in the diagonal are written to represent a mutational pathway leading to a sequence that is the closest to the WT variant, therefore, if we consider a pair of sequences 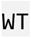 and 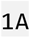, the diagonal consists of one mutation – 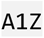, which would generate 1D hyperrectangle written as 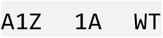 (Figure 3). If we consider two sequences which are more than one mutation away from each other, such as 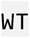 and 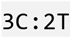, then the diagonal would be 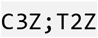 separated by a semicolon as part of a composite mutational path; this would be written as the following 1D hyperrectangle: 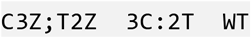. Higher dimensions are created by performing a pairwise comparison of hypercubes from lower dimensions and adding them together if they have a matching diagonal. For example, let us compare two 1D hypercubes – 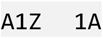 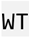 and 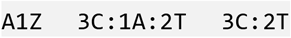 – given in Figure 3. Since they have matching diagonals, they can be combined into a 2D hyperrectangle 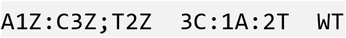 (Figure 3).

**Table 1.** A combinatorially complete dataset with mutations in K = 3 positions.

| Sequence* (mutations) | Fitness |
| --- | --- |
| WT | 0.7 |
| 3C:2T | 0.5 |
| 3C:1A:2T | 0.5 |
| 3C | 0.2 |
| 1A:2T | 0.7 |
| 1A | 0.9 |
| 3C:1A | 0.6 |
| 2T | 0.1 |

\*Sequences are represented as mutations added to the WT (e.g. 0C, mutation in position 0 to amino acid C). Letters A, T, C denote amino acids; letter Z – residue variant belonging to the WT sequence. Fitness values for each are randomly-generated numbers.

**Figure 3.**
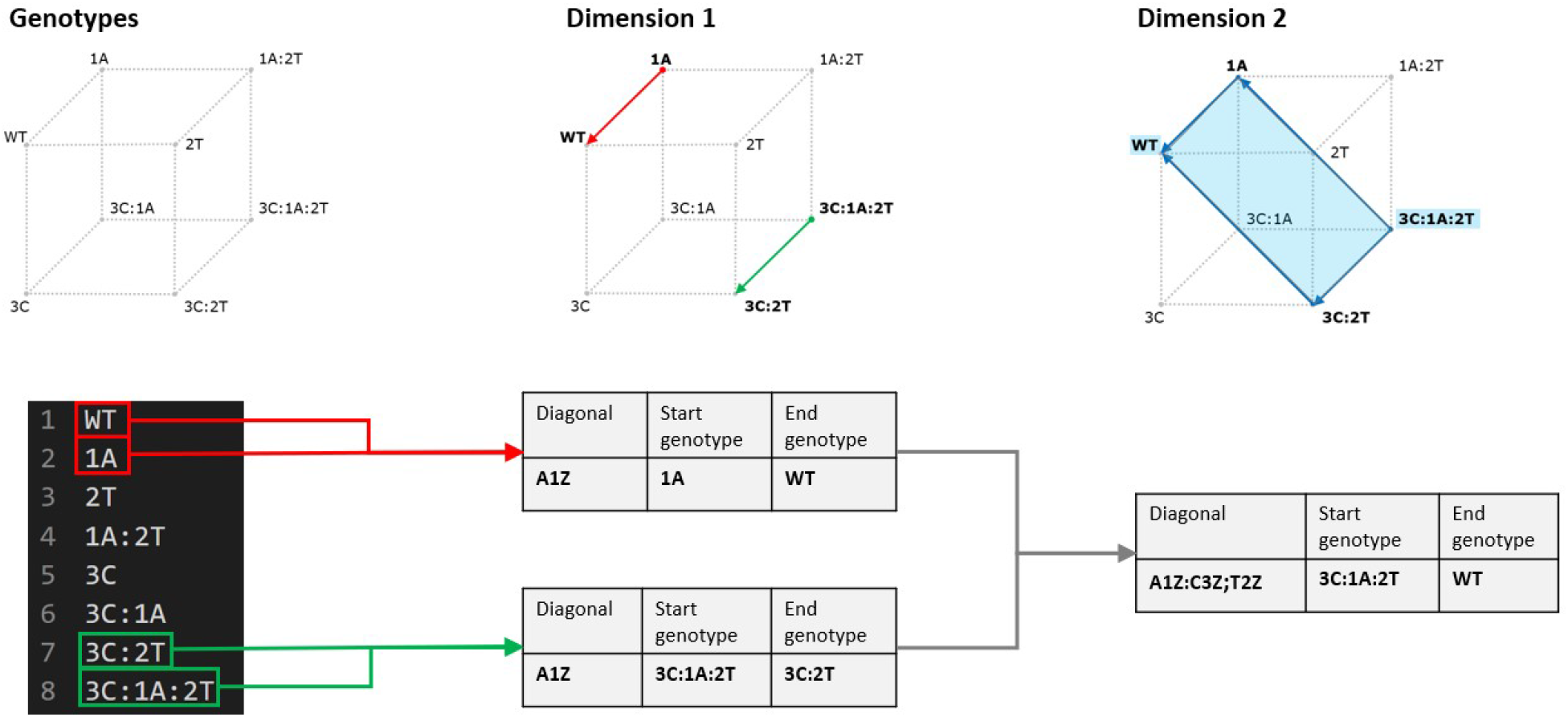
An example of constructing one 2D hyperrectangle 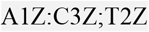 from a combinatorially complete dataset of eight sequences. Following (Esteban et *al*., 2019) Z represents a WT variant of an amino acid. Sequences are represented as mutations added to the WT (e.g. 3C, mutation in position 3 to amino acid C). Letters A, T, C denote amino acids; letter Z – residue variant belonging to the WT sequence.

The program terminates when, having made all pairwise comparisons of hypercuboids in a dimension, there are no matching diagonals.

### 3.2. Composite mutations dramatically increase the library of fitness landscape datasets

#### 3.2.1. Combinatorially complete datasets

Experimentally obtained fitness landscapes can have varying degrees of combinatorial completeness – a dataset might be lacking some of all possible combinations of the positions being considered. For this reason, we would expect to find different amounts of hypercubes and hyperrectangles depending on how combinatorially complete a given dataset is. Therefore, it is necessary to start the comparison of hypercubes and hyperrectangles by analyzing theoretically constructed combinatorially complete datasets of varying K (number of mutational positions under consideration).

For a combinatorially complete dataset, the numbers of hypercubes and hyperrectangles are constant for each K; accordingly, we can devise a mathematical equation for calculating the above structures. For a given K, we can calculate N (number of sequences or 0D hypercubes), determine the number of dimensions, and calculate the numbers of hypercubes, hyperrectangles, and both (hypercuboids) for each dimension and in total. Hypercubes follow a certain pattern of growth with each dimension. If we consider 1D hypercubes of a certain K, for example K = 2, we notice that increasing dimensionality by one (K = 3) yields the doubling of the lower-dimension edges (highlighted blue) as well as creation of the new edges connecting the two duplicate structures (highlighted red) (Figure 4A). Additionally, it can be noted that the number of the connecting edges increases by a factor of two with each K: going from K = 2 to K = 3, there are 4 new edges connecting the two duplicate planes (Figure 4A), while for the K = 3 to K = 4 transition there are 8 new edges (Figure 4B).

**Figure 4.**
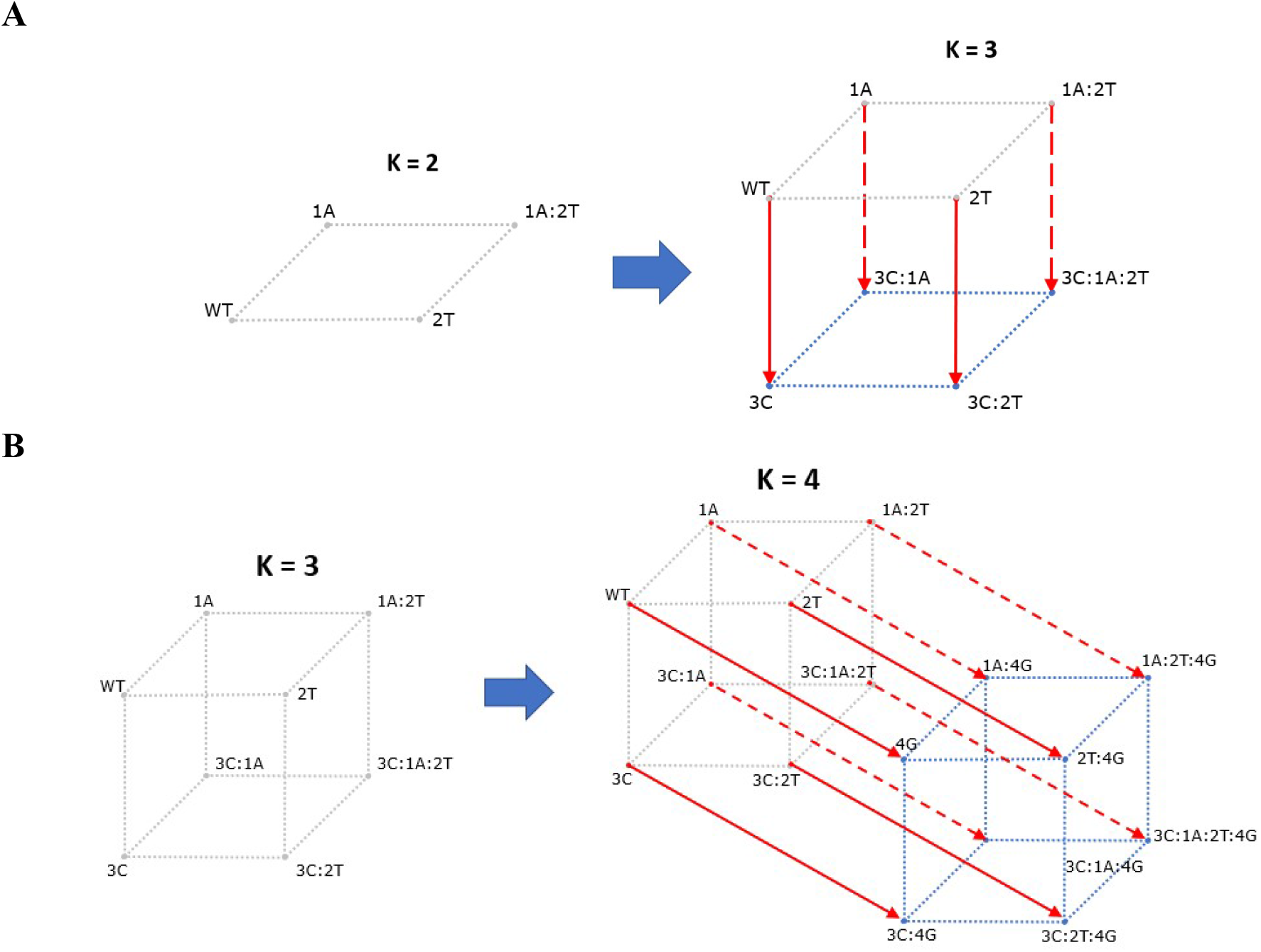
Procedure for constructing higher dimensional combinatorially complete space of K + 1 from a smaller dataset of K. **A**) Constructing K = 3 dataset from K = 2; **B**) Constructing K = 4 dataset from K = 3. Blue lines represent new edges duplicated from the previous dimension; red lines – new edges connecting the two duplicates.

From the patterns highlighted above, we can derive an equation sets that describe the growth of hypercubes, hyperrectangles, and hypercuboids for each dimension with each K.

##### Notes on notation

Let **K** – number of amino acid positions in a combinatorially complete dataset, **d** – dimension in that dataset, **x** – number of hypercubes, **y** – number of hyperrectangles, **z** – number of hypercuboids. General notation of number of hypercubes, hyperrectangles, and hypercuboids, respectively, is: 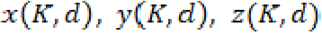. For a given *K*≥1, possible dimensions for the hypercuboids are integers belonging to a closed interval of *d* = [1, … *K*] (for hypercubes and hypercuboids), *d* = [1, …, *K* – 1] (for hyperrectangles). Using these definitions, we can generate number of hypercubes, hyperrectangles, and hypercuboids for each dimension for a given K:

Equation for calculating hypercubes:

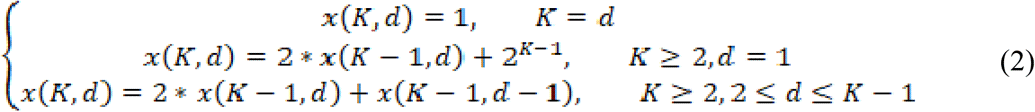

Equation for calculating hypercuboids:

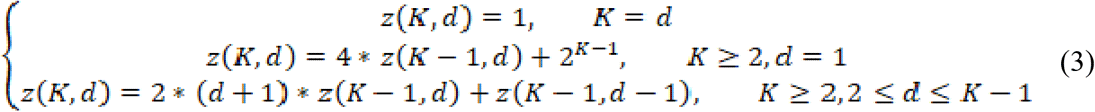

Equation for calculating hyperrectangles:

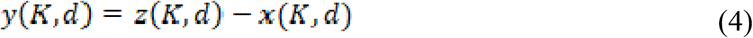

The mathematical formulae described above were implemented in Python, followed by the calculation of hypercubes and hyperrectangles for each K in the range of *K* = [1,26] (Figure 5).

**Figure 5.**
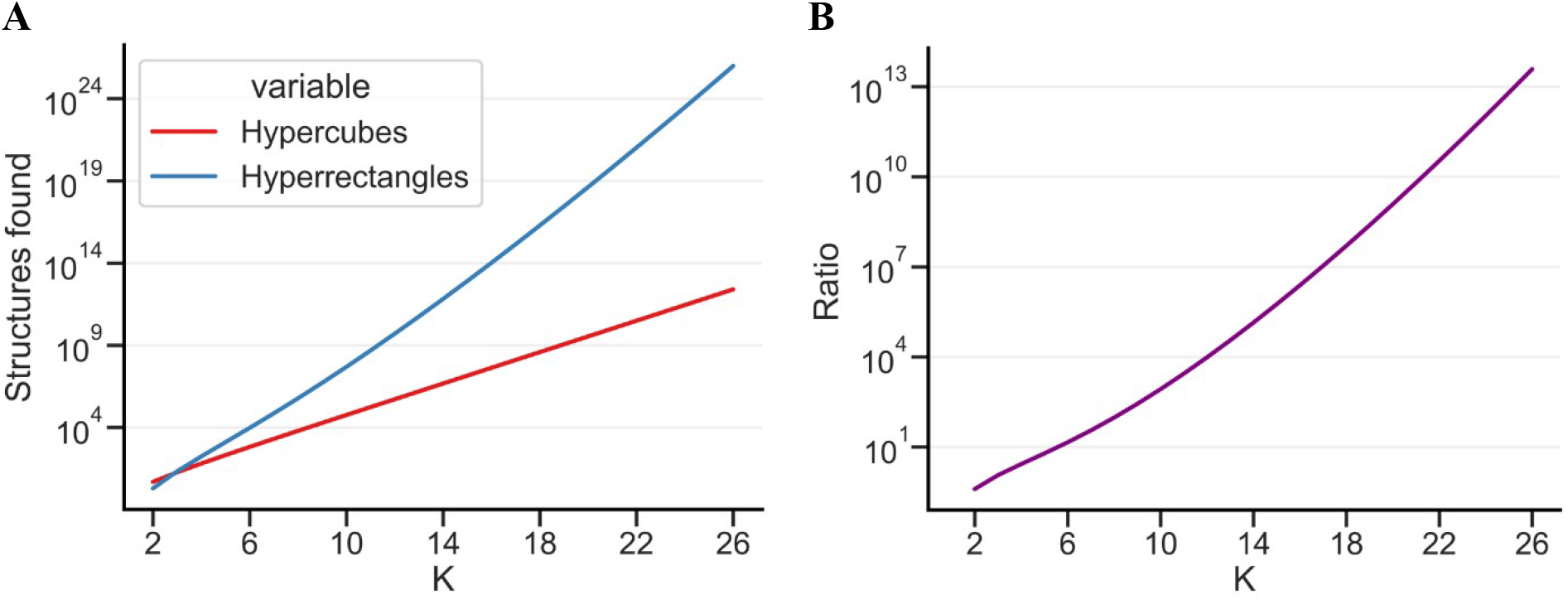
Numbers of hypercubes and hyperrectangles that were theoretically calculated for combinatorially complete datasets with K mutational positions. The lowest K calculated is 2 because K = 0 represents a WT sequence with no mutations being considered and at K = 1 there are no hyperrectangles found. **A**) Hypercubes and hyperrectangles theoretically calculated for each K; **B**) Ratio of hyperrectangles to hypercubes for each K.

Figure 5A shows that both hypercubes and hyperrectangles experience an exponential increase in numbers with increasing K. Nevertheless, hyperrectangles have a much higher rate of increase per K; this is evidenced in Figure 5B, where the ratio of hyperrectangles to hypercubes continuously increases with rising K, reaching a ratio as high as 10^13^ at K = 26. This demonstrates that in combinatorially complete datasets, there are many more hyperrectangles than hypercubes, providing a researcher with more data to investigate epistasis. The exact numbers for the Figure 5 can be seen in Supplement Table S1.

#### 3.2.2. Combinatorially incomplete datasets

Having completed the investigation of combinatorially complete datasets, we shifted our attention to analyzing large fitness landscape datasets. We analyze here the following datasets: green fluorescent protein (GFP) (Sarkisyan et al., 2016); segments 1-12 (S1-S12) of the protein IGPD (imidazoleglycerol-phosphate dehydratase) from His3 gene required for synthesis of histidine (Pokusaeva et al., 2019); the RNA recognition motif (RRM2) of *S. cerevisiae* poly(A)-binding protein (Melamed et al., 2013); WW domain of the protein hYAP65 (Araya et al., 2012).

Figure 6A shows the number of hypercubes and hyperrectangles for sample sizes of 100 – 1000 sequences. Figure 6B shows that the ratio of hyperrectangles to hypercubes is between 4 and 5 orders of magnitude. This is a much greater number than that for combinatorially complete theoretical datasets described in Chapter 3.2.1, where for K = 10 (equivalent to 1024 sequences) the ratio was 845, or just three orders of magnitude.

**Figure 6.**
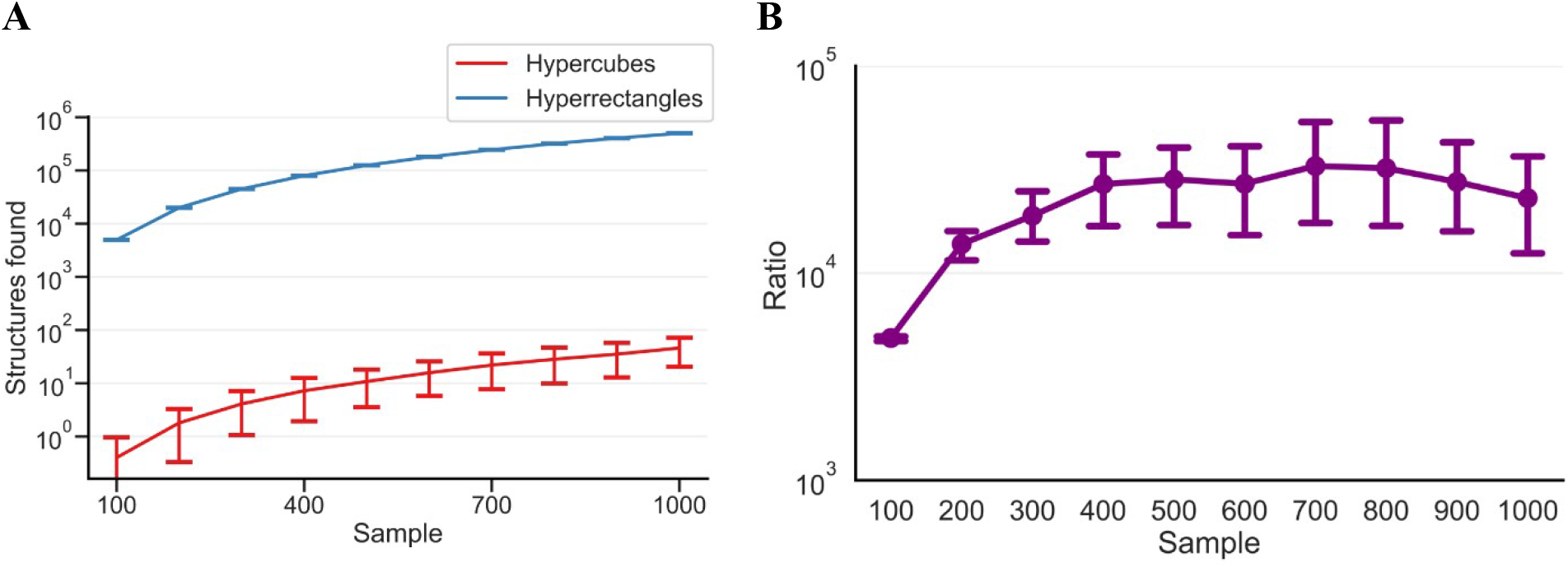
Comparison of hypercubes and hyperrectangles calculated for randomly-sampled fractions (100 – 1000 sequences; 10 replicates for each fraction) of different fitness landscapes (RRM2, WW, GFP, IGPDS1, IGPDS2, IGPDS3). Error bars represent SD for median values obtained for each dataset. **A**) Line plot of the amount of hypercubes and hyperrectangles per each sample size; **B**) Ratio of hyperrectangles to hypercubes.

### 3.3. Composite mutations allow detecting types of epistasis additional to that found by hypercubes

Using hyperrectangles for certain datasets can reveal additional types of epistasis not revealed by hypercubes. This is the case with the combinatorial complete subset from (Khan et al., 2011) displays example of ‘diminishing returns’ effects of beneficial mutations on fitness, or negative epistasis, which can explain the phenomenon of the diminishing fitness gain during adaptation (Chou et al., 2011). Let us consider the 3D combinatorially complete subspace comprising three positions (denoted in the article as **g**, **r**, and **p**), which represents a hypercube of K = 3; fitness values were extracted from Table S2 of (Khan et al., 2011). If within this subspace we consider 2D hypercubes (Figure 7A), we will only find negative epistasis (Figure 7B). However, when considering 2D hyperrectangles, there is a combination of datasets, specifically **p-r-gp-gr** (Figure 7C) which contains sign epistasis. Hence, the combinatorial subset of a real fitness landscape shows that hyperrectangles could find more sign epistasis as compared to hypercubes.

**Figure 7.**
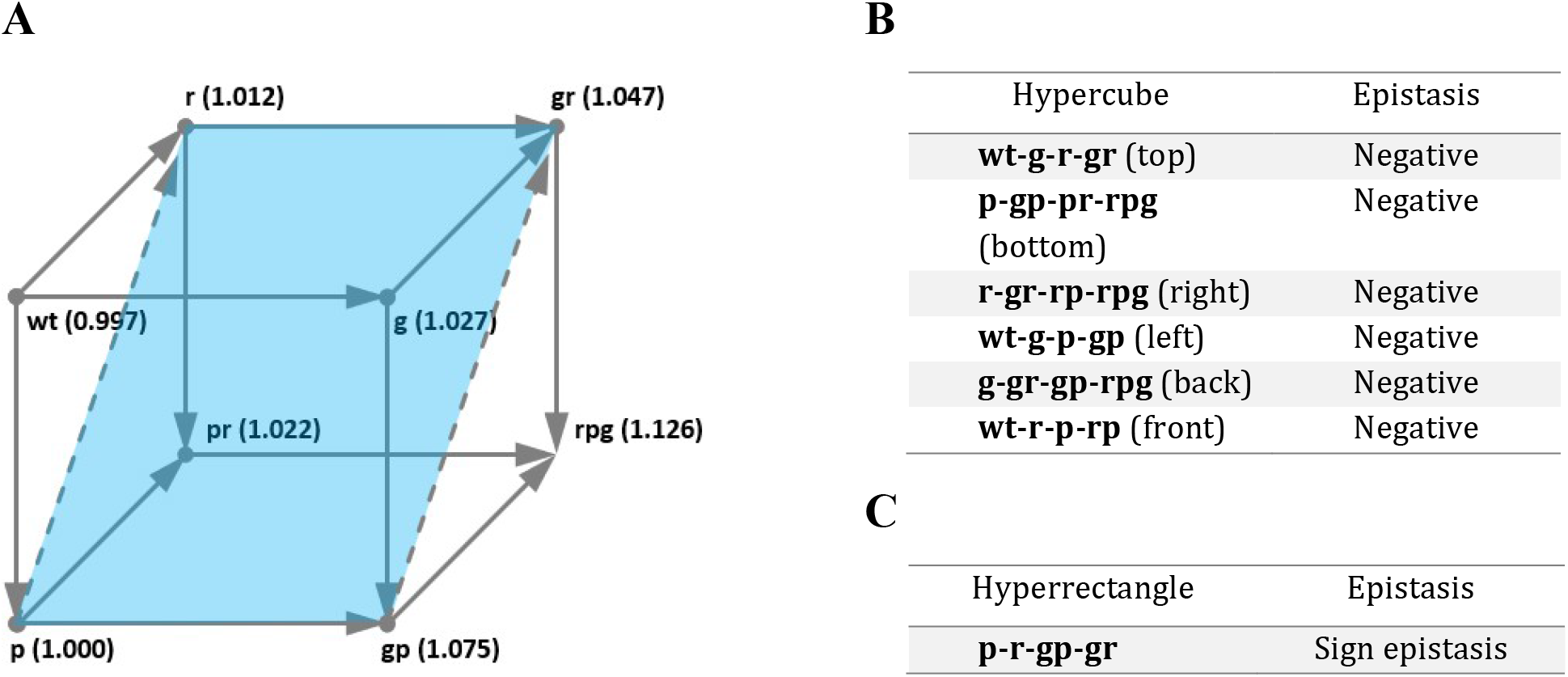
An example of an experimenal fitness landscape where hyperrectangles allow discovery of a type of epistasis not detected by hypercubes. **A**) Combinatorially complete subset (K = 3) of the dataset from the experimental fitness landscape (Khan et al., 2011). Solid lines represent all edges with H = 1, dashed lines = some of the edges with H > 1; **B**) List of 2D hypercubes found in the subset; **C**) An example of a 2D hyperrectangle (highlighted blue) found in the subset.

An intriguing example from Figure 7 warrants a more thorough investigation of the difference in types of epistasis between hypercubes and hyperrectangles – do hyperrectangles have significantly more percentage of a particular type of epistasis (negative, positive, sign, or reciprocal sign) compared to hypercubes? To conduct such an investigation, we analysed samples from datasets containing N ≥ 200 sequences by estimating types of epistasis in each hypercube and hyperrectangle for a dataset, then calculating percentage of each type of epistasis in each dataset, and finally performing Mann-Whitney U test to compare types of epistasis of the two groups (hypercubes and hyperrectangles) (Table 2).

**Table 2.**
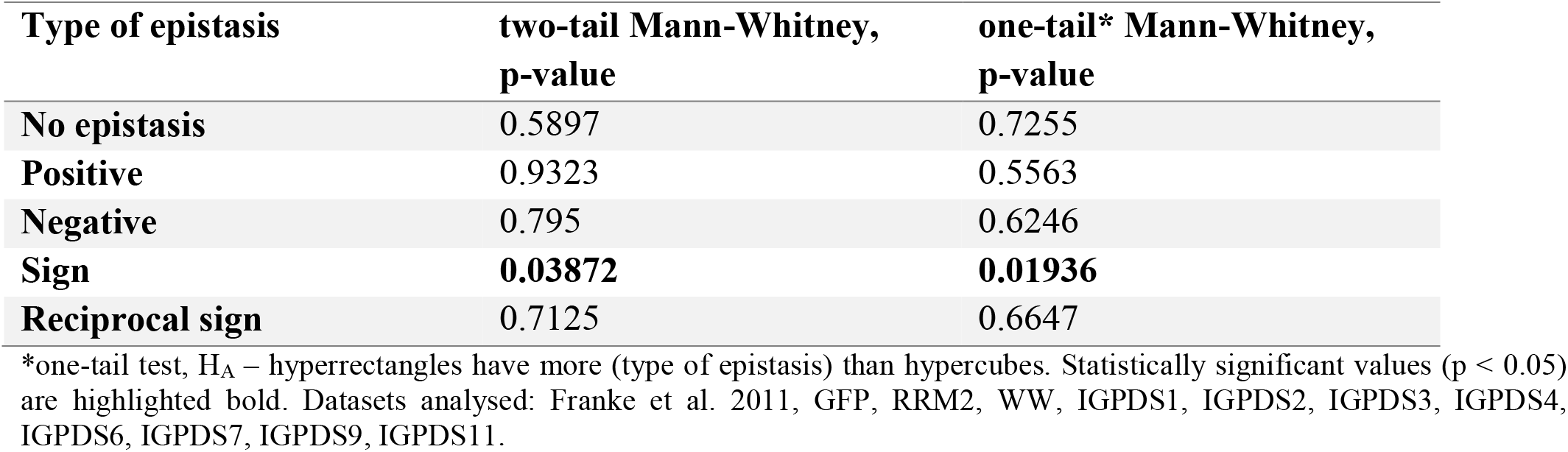
Difference in various types of epistasis (no epistasis, positive, negative, sign, and reciprocal sign) between hypercubes and hyperrectangles. Statistical test used is Mann-Whitney U test; for each type of epistasis, a single observation is a median of percentage of that type of epistasis for one dataset.

Table 2 shows that there is statistically significant difference between the two structures only for sign epistasis; furthermore, one-tail test indicates that hyperrectangles have more sign epistasis than hypercubes. Figure 8 shows the plot for residuals, where for each type of epistasis, we subtracted the percentage of one type of epistasis in hypercubes from hyperrectangles; each dot represents this action performed for each dataset. Indeed, the difference in percent of sign epistasis between the two structures averages around 5-10%, with one of the values reaching almost as high as 40% (Figure 8).

**Figure 8.**
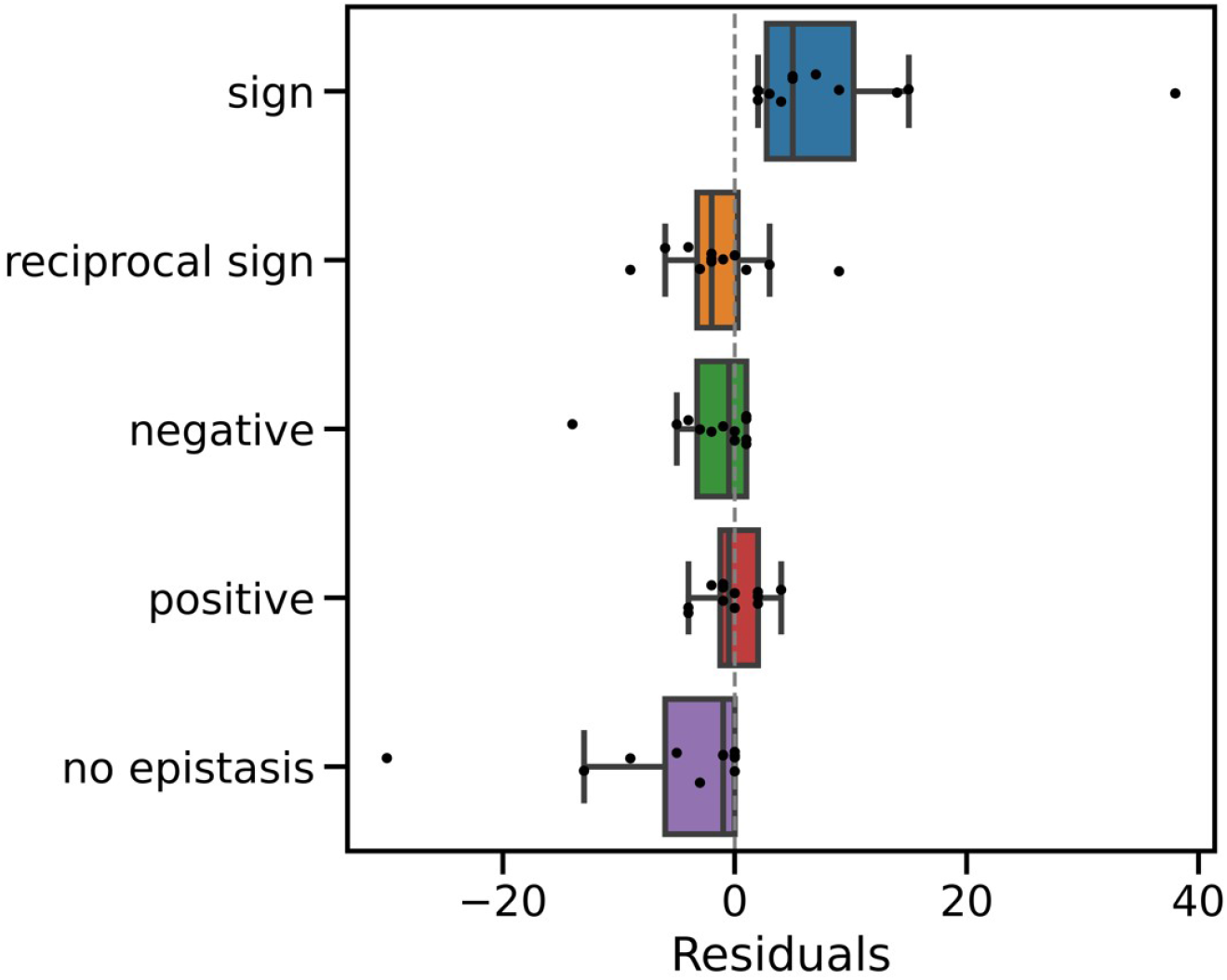
Distributions of residuals by types of epistasis. Residuals are calculated as the difference in percentage of a specific epistasis type in hyperrectangles and hypercubes for a specific dataset; each data point – one dataset. Horisontal line where Residuals = 0 represents no difference in percent of epistasis types between the two structures.

Therefore, we can see that hyperrectangles allow finding a higher percentage of sign epistasis.

### 3.4. Composite mutations allow for epistatic analysis in sparse datasets

If hyperrectange analysis gives more epistatic coefficients than hypercube one, there is a question: can we imagine a situation when the hypercube analysis does not give any epistatic coefficients at all while hyperrectangle analysis gives some epistatic coefficients? To explore this question, we took the first 1000 genotypes from some fitness landscapes, e.g. IGPDS2, IGPDS6, and IGPDS12, all subsets being combinatorially incomplete. Having calculated hypercuboids, we see that no hypercubes could be found, while hyperrectangles were found plentifully. This shows that there can be cases where using hyperrectangles can provide an information about some landsdcape where no hypercubes could be found. More datasets might be found in literature where researchers did not accomplish epistatic analysis because of absence of combinatorially complete datasets.

| Database | Hypercubes (dim=2) | Hyperrectangles (dim=2) |
| --- | --- | --- |
| IGPDS2 | 0 | 267 |
| IGPDS6 | 0 | 564 |
| IGPDS12 | 0 | 1,357 |

### 3.5. Comparison of epistatic coefficients obtained from hypercubes and hyperrectangles

In this section, we statistically compare epistatic coefficients *a*_12_ and their squares (*a*_12_)^2^ calculated for hypercubes and hyperrectangles. We would consider only those datasets which contain enough mutations and sequences to construct a combinatorially complete sample of K = 10, which is represented by *N* = 2^*K*^ = 2^10^ = 1024 sequences. The sampling of the datasets is performed because the combinatorial incompleteness of the datasets is likely to bias the results towards more epistasis in hyperrectangles as well as making the analysis of whole datasets too computationally intensive.

Of the large datasets, only four contained enough sequences to construct combinatorially complete datasets of K = 10 (Table 3). We used median as a measure of fitness, because the distributions are nonnormal with a high degree of skewness.

**Table 3.** Description of datasets used for calculating epistatic coefficients. For each dataset, we consider K = 10 mutational positions (N = 1024 sequences); consequently, for each dataset the numbers of 2D structures are the same: 11520 hypercubes and 7284736 hyperrectangles.

| Sample | Mutations | Fitness value<br>(median ± SD) |
| --- | --- | --- |
| IGPDS2 | A138P:F139Y:A140S:F141V:I142T:D143N:C156S:I159V:P160T:I162V | $0.81 \pm 0.44$ |
| IGPDS3 | F146L:K147Q:K150M:I151V:E153Q:L163M:D164H:A167S:G168R:G169S | $0.98 \pm 0.29$ |
| IGPDS4 | A170S:A171G:I172V:I174L:D177N:I179L:A194S:F195L:I197L:I199L | $1.00 \pm 0.079$ |
| IGPDS6 | A96G:I97L:S98T:K101Q:H104N:F116Y:S118Y:A119G:F120Y:A121C | $0.82 \pm 0.30$ |

Figure 9 shows the distribution of epistatic coefficients calculated for hypercubes and hyperrectangles. These distributions were shown to be non-parametric by the Anderson-Darling normality test; therefore, we used Mann-Whitney statistical test to compare the distributions of hypercubes and hyperrectangles in each dataset.

**Figure 9.**
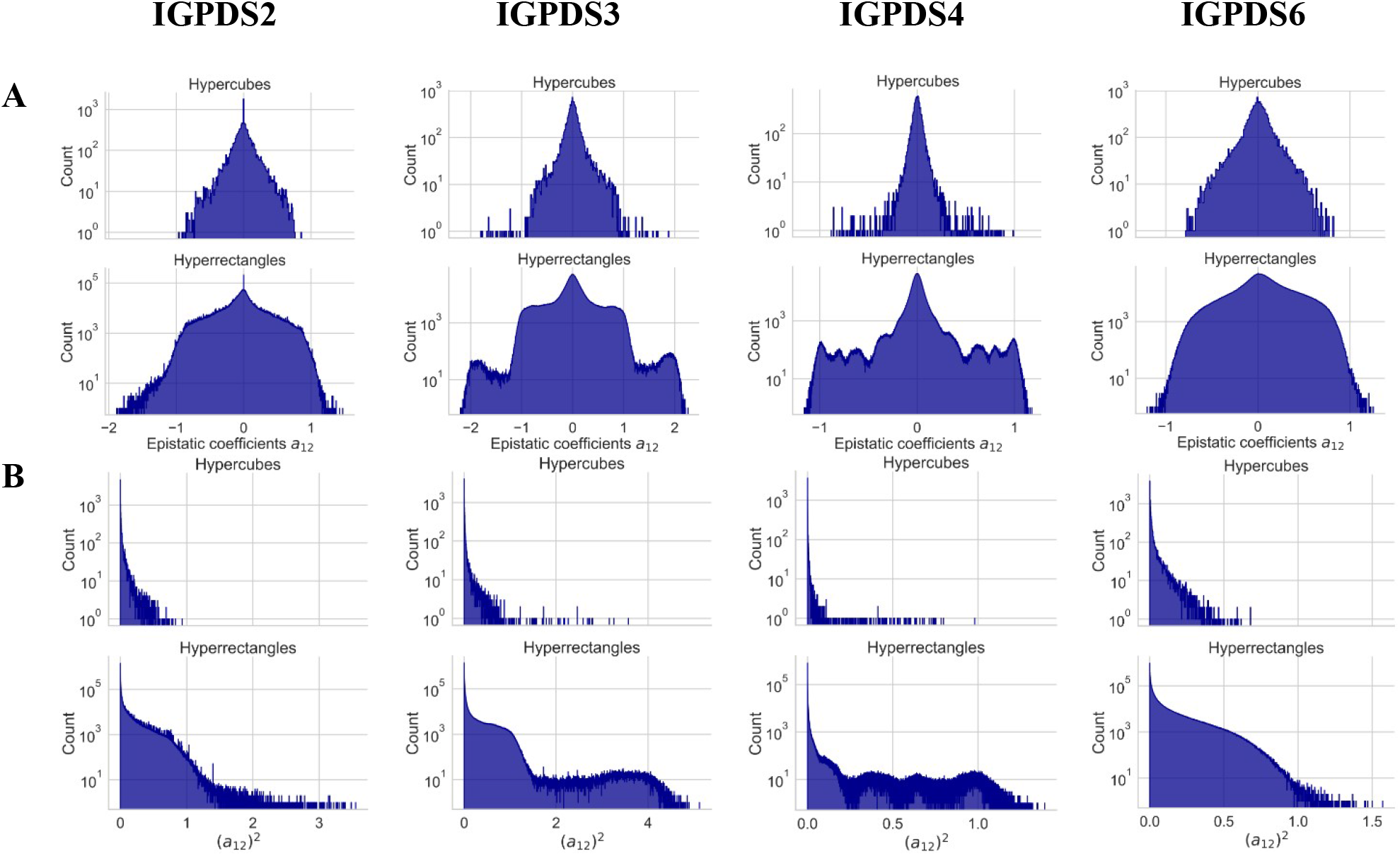
Distributions of epistatic coefficients for hypercubes and hyperrectangles. A) Histogram of epistatic coefficients *a*_12_; B) Histogram of squared epistatic coefficients (*a*_12_)^2^. Both A) and B) have logarithmic y-axes.

Mann-Whitney U test for *a*_12_ (two-tail) shows highly statistically significant (p < 0.001) difference in the distributions of hypercubes and hyperrectangles for IGPDS2, IGPDS3, and IGPDS6 (Table 4). To further confirm this difference, we conduct a one-tail Mann-Whitney U test for the squared epistatic coefficients (*a*_12_)^2^; squaring the coefficients exacerbates deviation from the null, or no epistasis. For these coefficients, we see that for all four datasets there is statistically significant difference (p < 0.001). Therefore, we can conclude that for the samples under consideration, hyperrectangles analysis give more epistasis than hypercubes one.

**Table 4.** Descriptions of statistical tests of hypercubes and hyperrectangles.

| Dataset | p-value for two-tailed<br>Mann-Whitney test, $a_{12}$ | Median ± SD of $(a_{12})^2$ | | p-value for one-tailed<br>Mann-Whitney test, $(a_{12})^2$ |
| --- | --- | --- | --- | --- |
|  |  | Hypercubes | Hyperrectangles |  |
| IGPDS2 | $p < 2.2e-16$ | $0.0045 \pm 0.074$ | $0.016 \pm 0.17$ | $p < 2.2e-16$ |
| IGPDS3 | $p = 9.067e-05$ | $0.0057 \pm 0.16$ | $0.018 \pm 0.30$ | $p < 2.2e-16$ |
| IGPDS4 | $p = 0.7743$ | $0.00074 \pm 0.047$ | $0.0011 \pm 0.10$ | $p < 2.2e-16$ |
| IGPDS6 | $p < 2.2e-16$ | $0.0055 \pm 0.062$ | $0.023 \pm 0.13$ | $p < 2.2e-16$ |

Mann-Whitney U test (two-tailed test), with the statistic value (W) and p-value. Alternative hypothesis: true location shift is not equal to 0 (two-tail test); Median and SD for distribution of epistatic coefficient squared $(a_{12})^2$ ; Mann-Whitney U test (one-tail test), with the statistic value (W) and p-value. Alternative hypothesis: true location shift is less than 0 (one-tail test). Median and SD are given for $(a_{12})^2$ and not $a_{12}$ , because for $a_{12}$ centers are at the 0 (no epistasis); for $(a_{12})^2$ , median represents deviation from the null.

Next interesting question to study is how the amount of epistasis measured by (*a*_12_)^2^ changes with increasing hypercuboid size. In other words, with increasing size of a hypercuboid (which is two for 2D hypercubes and 3+ for 2D hyperrectangles), would the epistasis increase, decrease, or no correlation would be found between the two variables? Figure 10 illustrates this pattern.

**Figure 10.**
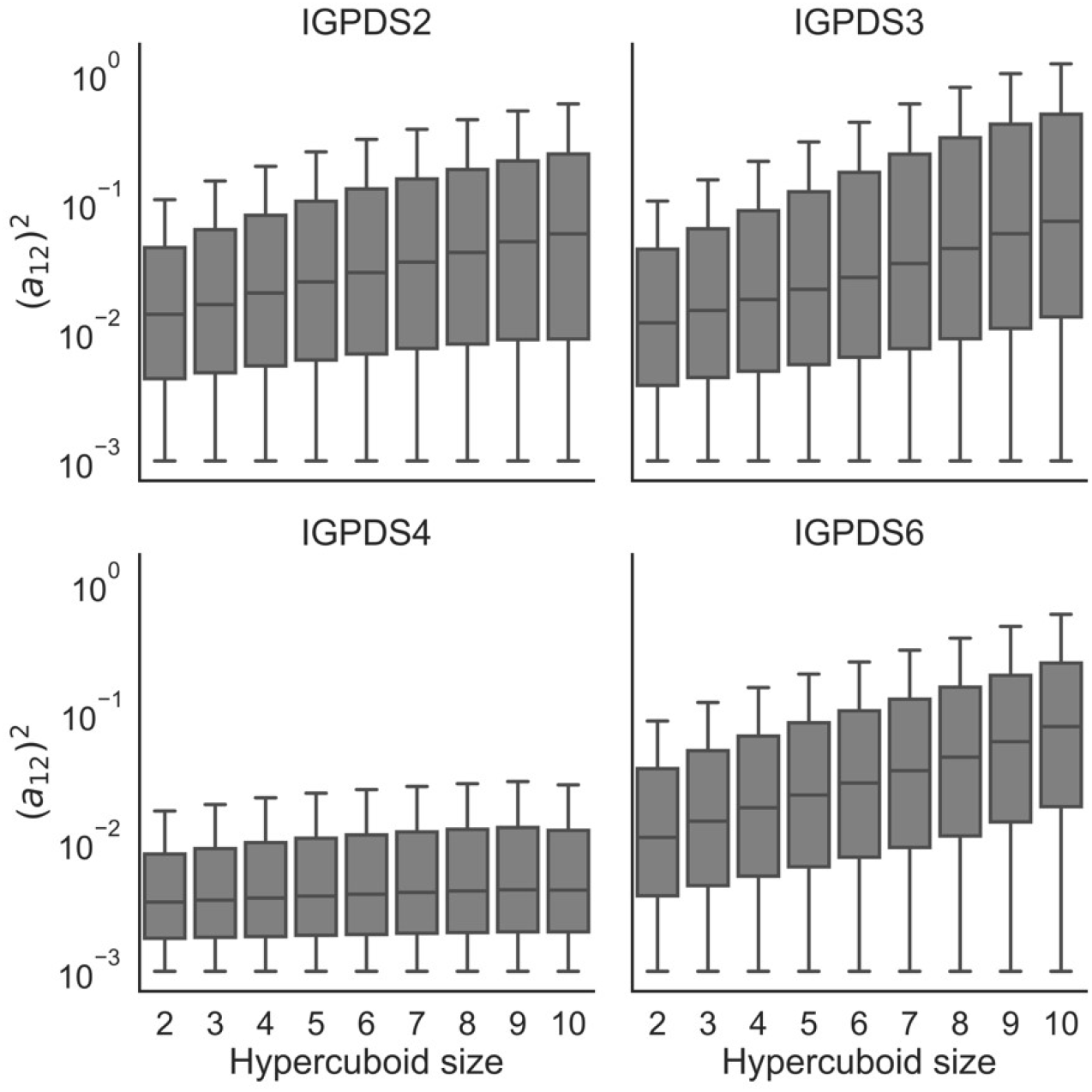
Dependence of epistasis and hypercuboid size. Epistatic coefficients (*a*_12_)^2^ < 10^−3^ were excluded from the analysis. Horisontal black bars in each box plot represent median values.

Figure 10 shows that there is a very big overlap between the individual boxes for different hypercuboid sizes; therefore, it is unlikely to be statistical difference in epistasis with the increasing hypercuboid size. Nevertheless, we can still see a trend of median value (depicted as black horizontal bar in each box subplot) increasing with hypercuboids size in datasets IGPDS2, IGPDS3, and IGPDS6.

### 3.6. Modelling hypercuboids after random removal of sample fractions

Having calculated the hyperrectangles / hypercubes ratio both for theoretically constructed combinatorially complete and experimental combinatorially incomplete datasets, we saw that the latter has a higher ratio – between 4-5 orders of magnitude (Section 3.2.2) compared to less than 3 orders of magnitude (Section 3.2.1). Therefore, in the present section we would investigate the dependence of ratio of hyperrectangles to hypercubes on the percentage of the dataset removed from analysis. In other words, if we move from analysing a combinatorially complete dataset to subsets with varying amounts of randomly removed sequences, in what direction would we see the ratio of hyperrectangles to hypercubes change? Here we calculated the ratios for *K* = [3,8]; we limited our analysis to K = 8 due to technical limitations by computational resources.

Figure 11 shows that in smaller fitness landscapes (K = 3, K = 4), the ratio tends to slightly increase. However, in a larger dataset of K = 8, we see that ratio decreased from nearly 100 down to around 40. The ratio of 100 for the whole combinatorially complete dataset at K = 8 (Figure 10) coincides with the calculation provided in Table S2; this is expected, as in both cases the dataset is combinatorially complete. The surprising observation comes as we start to remove sequences from the dataset. In case of K = 8, the ratio goes down, eventually reaching the level below 40.

**Figure 11.**
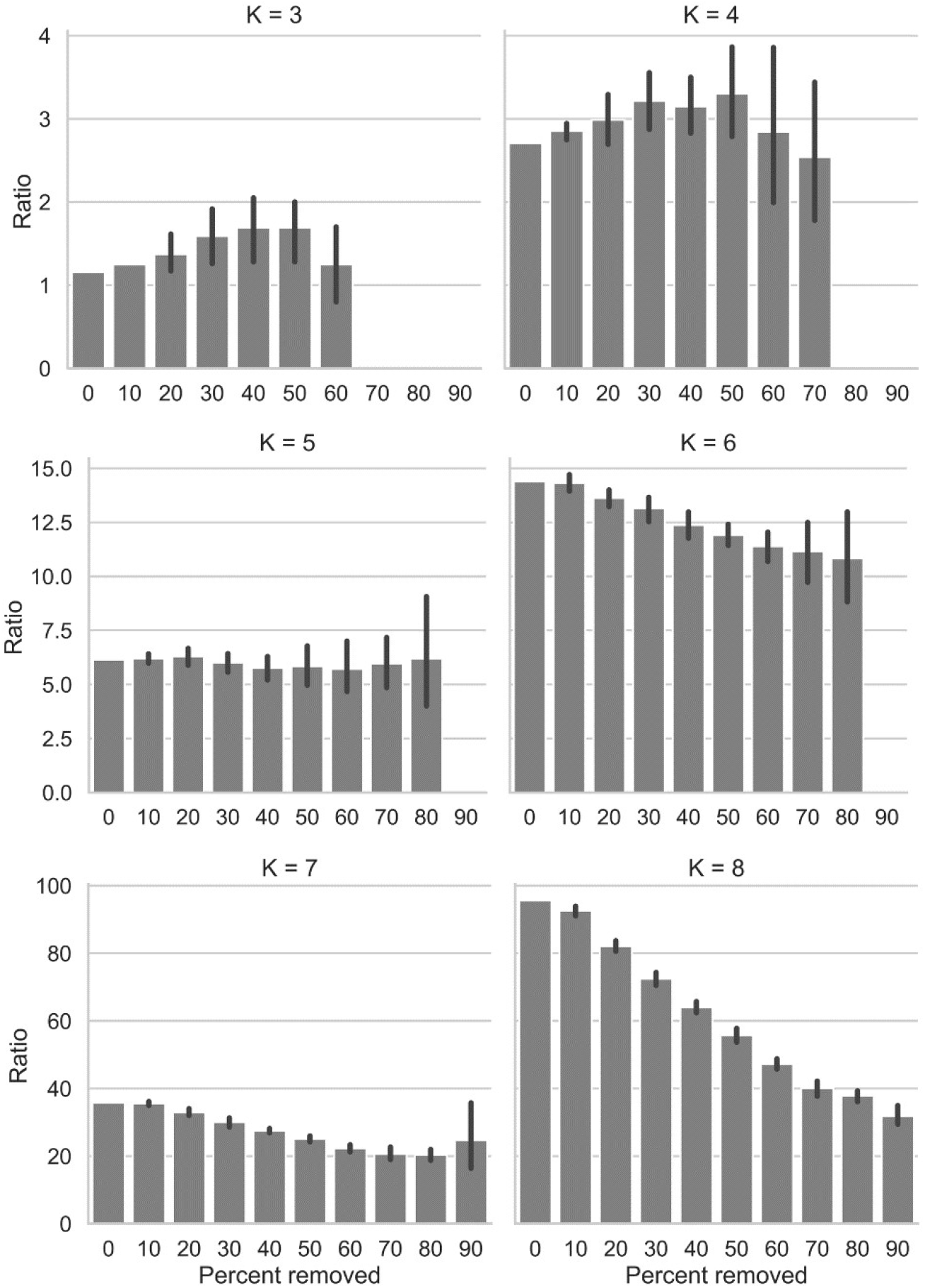
Ratios of hyperrectangles to hypercubes for increasing percentage of removed sequences (0%, combinatorially complete dataset; 90%, starting dataset with 90% of its sequences randomly removed), stratified by the number of positions K under consideration (*K* = [3,8]).

## 4. Discussion

Despite numerous articles on the topic, epistasis remains a cryptic phenomenon. Its understanding influences our comprehension of fitness landscapes, as epistasis is a primary determinant of the shape of a fitness landscape by influencing its parameters such as ruggedness and curvature (Beerenwinkel et al., 2007). Being considered from the standpoint of evolution, fitness landscapes are traversed by means of single mutations, with the evolutionary step being equal to 0.5-1 mutation per generation in eukaryotes (Barrick and Lenski, 2013). Nonetheless, there are some natural genetic phenomena including multiple mutait<u>i</u>ons; these include recombination and transposition. Accordingly, studying many substitutions at a time could also be relevant even for the process of evolution; however, we understand that finding hyperrectangle structures in evolutionary data has practically zero chances.

Previously, there have been limited attempts to implicitly utilize composite mutations to study epistasis. Beerenwinkel with colleagues (Beerenwinkel et al., 2007) used a so-called “new geometric approach”, where “the geometric framework allows not only tests of the standard pairwise interactions but also non-pairwise tests that gave new insights into (i) the relationship between the form of epistasis and the individual mutational effects, and (ii) variation between mutations in their mixing ability with other mutations”. In their analysis, they were able to show that using non-standard tests over the standard ones enabled them to find more epistasis in the fitness landscape.

Other research groups have studied effects of mutations in different contexts. Pokusaeva with colleagues, for example, explored effects of the same mutations in far genetic contexts (Pokusaeva et al., 2019). Thus, they implicitly studied 2D hyperrectangles where two edges have H = 1 while the other two edges can be of any length.

Table 2 (Section 3.3) showed some intriguing results. For most types of epistasis (positive, negative, reciprocal sign, or no epistasis), percentages between the two structures (hypercubes and hyperrectangles) do not seem to differ. At the same type, there is a difference in percentage of sign epistasis between the two groups (p = 0.03872), with hyperrectangles having a higher percentage of sign epistasis than hypercubes (p = 0.01936). This significant difference was observed even with such a small sample size of n = 12 datasets; therefore, we could expect that with a larger sample of datasets the statistical power may be even stronger and the difference – more statistically significant. Figure 6 shows that the difference in percent of sign epistasis between the two structures averages around 5-10%, with one of the values reaching almost as high as 40%.

It is also interesting to compare the results of the ratio of hyperrectangles to hypercubes for combinatorially complete datasets (Section 3.2.1), for real experimental datasets (Section 3.2.2), and for the random removal experiment (Section 3.6). For convenience, some of the results are summarized in Table 5. If we consider a dataset of K = 7 (corresponding to 128 sequences), a theoretical combinatorially complete dataset would have a ratio of 36. The ratio is the same in the random removal experiment before randomly removing any sequences; after 90% of such removal, the ratio goes down to 21. In general, if we consult Table 5B, C, we can see that after having removed 90% of sequences in a dataset, the ratio goes down dramatically. However, Table 5D shows the ratios that far exceed those for the other datasets.

**Table 5.** Comparison of the hyperrectangles / hypercubes ratio for combinatorially complete datasets, real combinatorially incomplete datasets, and experimentally modelled incomplete datasets, for varying sequences.

| Sequences (K) | A) Comb complete<br>ratio | B) Modelling<br>incompleteness:<br>0% removed | C) Modelling<br>incompleteness:<br>90% removed | D) Real<br>experimental<br>dataset |
| --- | --- | --- | --- | --- |
| 64 (K = 6) | 14 | 14 | 7 | - |
| 128 (K = 7) | 36 | 36 | 21 | 4867 |
| 256 (K = 8) | 96 | 96 | 31 | 18943 |
| 512 (K = 9) | 275 | - | - | 28360 |
| 1024 (K = 10) | 846 | - | - | 23020 |

values are rounded to the nearest whole integer. Ratios for “real experimental dataset” are given as those for the number of sequences closest to the corresponding K: for K = 7, 100 sequences; for K = 8, 300 sequences; for K = 9, 500; for K = 10, 1000.

Such a drastic difference in ratios could be explained by a systematic difference between incompleteness of the datasets given in Chapters 3.2.2 and 3.6. While in the latter case, the combinatorially incomplete datasets were created by removing fractions consisting of randomly sampled sequences, the former could have a bias for excluding mutations in a pattern which would lead to the increase in the ratio of hyperrectangles to hypercubes. Such a bias could be exemplified by choice of clusters of mutations for the experimental datasets that are separated spatially by many amino acids positions. In this case, hypercubes would only be found in the respective clusters of mutations that are close to each other, while hyperrectangles would be calculated within as well as between the clusters, leading to a great increase in numbers of hyperrectangles. This explanation could be tested in the future research as the dependence of ratio of hyperrectangles to hypercubes on the pattern of combinatorial incompleteness of the dataset.

Overall, we demonstrate here that epistatic analysis based on composite mutations and hyperrectangles not only produce more dramatically more data containing valuable information about epistasis but also allows to reveal new types of epistasis. We expect that the proposed approach will have a high importance for the analysis of the evolutionary landscapes published by now and in the future.

## Abbreviations

WT: wild type
GFP: green fluorescent protein
IGPD: imidazoleglycerol-phosphate dehydratase
IGPDSX: imidazoleglycerol-phosphate dehydratase, segment X
BASH: bourne again shell
H: hamming distance

