## Supplementary Material for "Composite mutations give an extra insight into epistasis"

### Supplementary information

**Table S1**. Numbers of hypercubes, number of hyperrectangles, and their ratio for $K = [1, 28]$.

| K | Sequences | Hypercubes | Hyperrectangles | Ratio |
|---|---|---|---|---|
| 1 | 2 | 1 | 0 | - |
| 2 | 4 | 5 | 2 | 0.4 |
| 3 | 8 | 19 | 22 | 1.16 |
| 4 | 16 | 65 | 176 | 2.71 |
| 5 | 32 | 211 | 1,296 | 6.14 |
| 6 | 64 | 665 | 9,570 | 14.39 |
| 7 | 128 | 2,059 | 73,718 | 35.80 |
| 8 | 256 | 6,305 | 602,880 | 95.62 |
| 9 | 512 | 19,171 | 5,264,768 | 274.62 |
| 10 | 1024 | 58,025 | 49,075,870 | 845.77 |
| 11 | 2048 | 175,099 | 486,849,800 | 2780.43 |
| 12 | 4096 | 527,345 | 5,120,374,000 | 9709.72 |
| 13 | 8192 | 1,586,131 | 5.68765e+10 | 35858.64 |
| 14 | 16384 | 4,766,585 | 6.649152e+11 | 139495.1 |
| 15 | 32768 | 14,316,140 | 8.155326e+12 | 569659.6 |
| 16 | 65536 | 42,981,180 | 1.046525e+14 | 2434844 |
| 17 | 131072 | 129,009,100 | 1.401573e+15 | 10,864,140 |
| 18 | 262144 | 387,158,300 | 1.954687e+16 | 50,488,060 |
| 19 | 524288 | 1,161,737,000 | 2.833149e+17 | 243,871,800 |
| 20 | 1048576 | 3,485,736,000 | 4.259998e+18 | 1,222,123,000 |
| 21 | 2097152 | 1.045826e+10 | 6.634162e+19 | 6,343,469,000 |
| 22 | 4194304 | 3.137687e+10 | 1.068441e+21 | 3.405187e+10 |
| 23 | 8388608 | 9.413479e+10 | 1.777082e+22 | 1.887806e+11 |
| 24 | 16777216 | 2.824128e+11 | 3.048665e+23 | 1.079507e+12 |
| 25 | 33554432 | 8.472551e+11 | 5.388303e+24 | 6.359718e+12 |
| 26 | 67108864 | 2.541799e+12 | 9.800905e+25 | 3.855893e+13 |
